# Hepatic GCGR is required for the superior weight loss effects of a structurally related analogue of the dual GCGR/GLP1R agonist survodutide

**DOI:** 10.1101/2024.09.09.611134

**Authors:** Fen Long, Manuel Klug, Tenagne D. Challa, Vissarion Efthymiou, Christian Wolfrum, Carla Horvath

## Abstract

The dual glucagon/glucagon-like peptide 1 receptor (GCGR/GLP1R) agonists have superior efficacy in promoting weight loss and metabolic improvements in obesity and metabolic dysfunction-associated steatohepatitis (MASH) than current available mono-agonists. However, the mechanisms underlying these benefits are not fully understood. While the effects on appetite regulation and glucose control through GLP1R agonism are well established, the role of GCGR agonism in promoting weight loss and metabolic changes is less defined. Using a dual GCGR/GLP1R agonist BI 456908 and a selective GLP1R agonist semaglutide, we could show that the dual agonist achieved superior weight loss efficacy by engaging hepatic GCGR without adversely affecting glucose control. Furthermore, we could demonstrate that hepatic GCGR is critical for facilitating plasma and liver lipid clearance stimulated by the dual agonist. Overall, these findings highlight the crucial metabolic contributions of hepatic GCGR to the efficacy of combined GCGR/GL1R activation.

## Introduction

Recent advancements in peptide-based anti-obesity drugs, comprising single and different combinations of dual or triple agonists, have revolutionized obesity treatment (1). The addition of the GCGR component to GLP1R agonism has improved weight loss efficacy compared to GLP1R stimulation only and may provide additional cardiometabolic benefits in diet-induced obese mice (2). GCG, the major counter-regulatory hormone of insulin, plays a critical role in systemic metabolic control and regulates multiple catabolic pathways affecting lipid, glucose and amino acid homeostasis (7). Despite these potential benefits, the use of GCG as a monotherapy is limited by its hyperglycaemic effects (8). The metabolic actions of GCG are primarily mediated by the direct activation of the GCGR, with the liver being the principal target organ due to its highest expression of GCGR (9).

Survodutide, a dual GCGR/GLP1R agonist, has demonstrated superior weight loss effects in individuals with obesity (currently in Phase III clinical trials), as well as significant improvements in metabolic-associated steatohepatitis (MASH) and fibrosis (3, 4). These outcomes suggest that GCGR agonism provides direct metabolic advantages such as increased energy expenditure (EE), lipolysis and mobilization of hepatic fat stores. However, the precise contribution of GCGR activation and the underlying target tissues remains unclear. In general, the potent effect of survodutide on body weight reduction can be attributed to the superior of actions of simultaneous GCGR and GLP1R activation. While GLP-1R activation primarily suppresses appetite, increases satiety and delays gastric emptying - highlighting its therapeutic efficacy for the treatment of obesity and type II diabetes, as exemplified by the clinical use of semaglutide. GCGR activation has been proposed to promote weight loss through increased EE (5). This is in line with observations that only survodutide but not semaglutide treatment increased EE in obese mice, and underscores the potential of GCGR activation on resting metabolic rate (2). Furthermore, this dual mode of action of GCGR/GLP1R agonists could potentially prevent declining EE that parallels weight loss (6).

In this study, we investigated the metabolic contribution of hepatic GCGR activation of a dual GCGR/GLP-1R agonist BI 456908, which exhibits structural and pharmacological similarities to survodutide (10).

## Results

BI 456908, alongside the GLP1R selective agonist semaglutide, were used to investigate the role of the GCGR component in the weight-lowering effects of dual GCGR/GLP1R agonists (Fig. 1a). To determine to which extent these effects were mediated by hepatic GCGR signalling, we utilized a liver-specific GCGR knockout (LKO) mouse model to differentiate the hepatic GCGR signalling component.

**Fig. 1.**
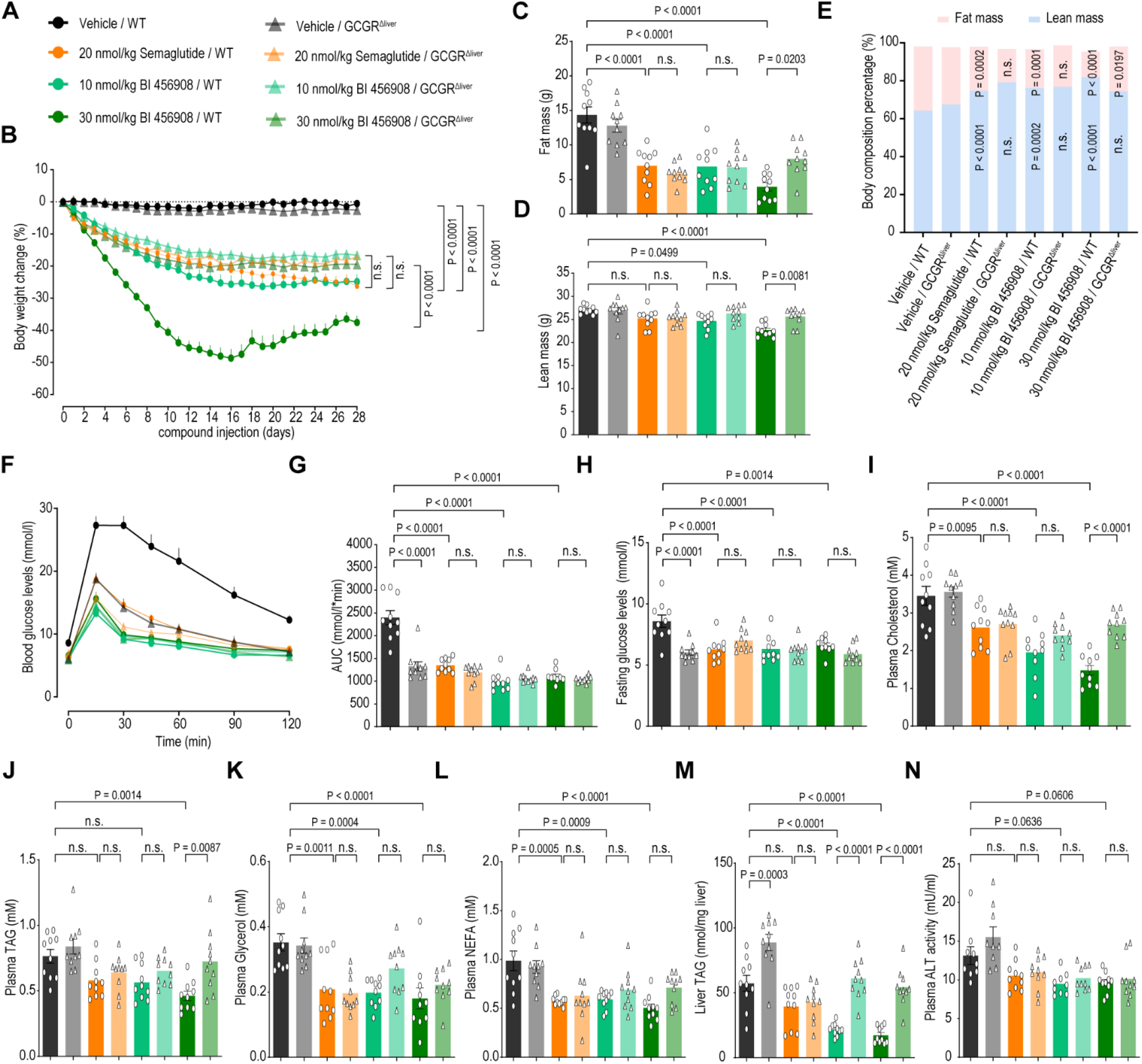
Metabolic improvements following 29 days of semaglutide or BI 456908 treatment in DIO WT and hepatic GCGR KO mice. (A)Group annotations. (B)Percentage change in body weight during the 29-day treatment with daily subcutaneous injection of either 20 nmol/kg semaglutide or 10/30 nmol/kg BI 456908. (n=10 per group). (C-E) Body fat mass (C), lean mass (D), and body composition proportions (E) at study termination (n=10 per group). (F-H) Intraperitoneal glucose tolerance test (ipGTT) (F) was conducted on day 21 of treatment. Mice were fasted for 4 hrs prior to glucose injection (2 g/kg). (G) Area under the curve for ipGTT and (H) Fasting blood glucose levels measured at baseline. (n=10 per group). (I-L) Plasma levels of cholesterol (I), TAG (J), glycerol (K), and NEFA (L) at study termination (n=10 for per group). (M)Liver TAG levels at study termination (n=10 for per group). (N)Plasma ALT activity at study termination (n=9 for Vehicle /GCGR^Δliver^ group and n=10 for other groups). Data presented as mean ± SEM. Each data point represents an individual animal. Statistical significance indicates comparison between treated-WT mice and WT controls or between treated-WT and treated-GCGR LKO mice. Statistical analysis was performed using two-way ANOVA with Tukey’s post hoc multiple comparison test (B-N).

As expected, semaglutide (20 nmol/kg) resulted in ∼20% weight loss in both diet-induced obese (DIO) wildtype (WT) and LKO mice following four weeks of treatment (Fig 1b). BI 456908 led to a dose-dependent weight loss in DIO WT mice. At 10 nmol/kg dose, WT mice lost ∼20 % weight, with small difference between WT and LKO mice. At 30 nmol/kg dose, WT mice showed a substantial ∼40 % weight loss, while LKO mice exhibited only marginally higher weight loss compared to the lower dose. Importantly, the pronounced weight loss efficacy was significantly diminished in LKO mice and approximated the effects of semaglutide, underscoring the critical role of hepatic GCGR activation in BI 456908’s efficacy and its additive effects to GLP1R agonism.

Weight loss induced by semaglutide and BI 456908 was mostly attributable to fat mass loss (Fig. 1C), although some lean mass loss was observed in BI 456908-treated WT mice at the high dose (Fig 1D). Both agonists altered body composition, decreasing fat mass and increasing lean mass proportions (Fig 1C). However, at the high dose, BI 456908 led to a more pronounced decrease in fat mass proportion in WT mice compared to LKO mice. These data highlight the importance of hepatic GCGR in the weight loss effects of BI 456908 and suggest a depletion of fat storage to fuel lipid utilization in the liver.

Next, we assessed whether BI 456908 could improve weight loss without compromising glycaemic control due to a potential induction of hepatic glucose production via GCGR. BI 456908 ameliorated glucose tolerance at both doses with comparable effects to semaglutide (Fig 1F-G). Notably, fasting blood glucose levels were decreased in both semaglutide and BI 456908 treated mice, independently of the genotype (Fig. 1H). These improvements were maintained in LKO mice, highlighting the effectiveness of the GLP1R component in restoring glycaemic control.

Both semaglutide and BI 456908 lowered plasma cholesterol, demonstrating GLP1R’s benefits in plasma lipid clearance (Fig. 1I). Hepatic GCGR activation provided additional cholesterol reduction, as evidenced by the differential effects of high-dose BI 456908 between WT and LKO mice. Furthermore, only high-dose BI 456908 significantly reduced plasma triacylglycerol (TAG) levels, an effect lost in LKO mice (Fig. 1J), indicating the metabolic advantages of hepatic GCGR activation. Both semaglutide and BI 456908 reduced plasma glycerol (Fig. 1K) and non-esterified fatty acid (NEFA, Fig 1L) levels to a comparable extent. Although low-dose BI 456908 did not outperform semaglutide in weight loss, it significantly decreased liver TAG levels via hepatic GCGR activation.

GCGR stimulation in the liver induces lipolysis via PKA signaling and promotes fatty acid oxidation. Thus, we quantified hepatic fat deposition to evaluate a potential improvement in MALFD following agonist treatments. As expected, LKO vehicle-treated mice displayed higher liver TAG content compared to WT-mice. In contrast to Semaglutide, which did not significantly reduce TAG load, low and high doses of BI 456908 enhanced liver TAG clearance in WT mice but were ineffective in LKO mice. In line with this, BI 456908 effectively reduced plasma ALT levels (Fig. 1 N), substantiating improved liver metabolic health, however these effects were independent of GCGR activation.

Collectively, these findings underline the essential role of hepatic GCGR in the weight loss efficacy of dual GCGR/GLP1R agonists and highlight additional metabolic benefits beyond weight reduction through hepatic GCGR activation.

## Discussion

The concurrent activation of GCGR and GLP1R signaling in different target tissues modulates both sides of the energy balance as reflected in reduced energy intake and elevated EE, which cumulates in robust weight loss (2). This implies that the weight loss effects of dual GCGR/GLP1R agonism are separable into a food intake dependent and independent component. The glucagon-induced rise in EE has been demonstrated across various species, including human and rodent models (11), but the responsible mechanisms are as of yet unknown. We show here that functional hepatic GCGR-signaling is required for the highly efficacious additive weight loss effect of BI 456908, suggesting that the food intake independent component translates to enhanced liver EE. Our data further indicate that the liver is an integral component of systemic EE and modulation of hepatic EE can influence whole-body energy homeostasis to a physiologically relevant extent. The precise mechanisms and contributors of liver-mediated EE are not fully elucidated, but hypoaminoacidemia and intact hepatic Farnesoid X receptor (FXR) signaling have been identified as prerequisites for the GCG-induced increase in EE (5, 12).

Increased EE is commonly accompanied by a compensatory rise in food intake to satisfy the caloric requirements (13), which might be limited in case of BI 456908 treatment due to the appetite suppressing effect of GLP1R agonism. Instead, the enhanced energetic demand is potentially fuelled by the depletion of adipose tissue lipid stores, as apparent by the more pronounced fat mass loss in BI 456908 compared to semaglutide treated mice. Furthermore, all treatment groups ameliorated various parameters of dyslipidaemia, substantiating the systemic metabolic benefits and therapeutic potential of dual GCGR/GLP1R agonists. This lipid-lowering effect might also stem from the global improvement in metabolic control linked to weight loss, the higher metabolic activity in the liver or direct GCG-controlled regulation of molecular processes. In line with the latter, BI 456908 induced a dose-dependent reduction in plasma cholesterol levels. This could be due to the regulation of low-density lipoprotein (LDL) cholesterol homeostasis by hepatic GCGR by promoting the degradation of proprotein convertase subtilisin/kexin type 9 (PCSK9), which in turn reduces LDL-receptor internalisation and accelerates LDL-C clearance from the circulation (14).

Survodutide, structurally closely related to BI 456908 and qualified for clinical trials (10), significantly reduces liver fat content and exerts beneficial effects on liver fibrosis in patients with confirmed disease stages (3). We could recapitulate a decrease in liver TAG accumulation in response to BI 456908 treatment, which was dependent on hepatic GCGR expression and corroborates the lipolytic activity of GCGR activation. It is well established that GCG triggers lipolysis as well as mitochondrial fatty acid oxidation in the liver via transcriptional-dependent and independent effects mediated by activated phospholipase C (PLC) and protein kinase A (PKA) downstream of GCGR (15). Importantly, Petersen et al. very recently demonstrated that a physiological elevation in circulating glucagon levels during a two hours infusion boosts hepatic mitochondrial oxidation in control subjects and MAFLD patients (16). At the molecular level, increased EE is intrinsically linked to higher mitochondrial respiratory rates and the utilization of reducing equivalents gained from substrate oxidation to generate a proton gradient across the inner mitochondrial membrane (17). The energy stored therein is used to power ATP-production through the ATP synthase complex or alternatively to produce heat via the uncoupling protein 1 (UCP1), which is uniquely expressed in brown and beige adipocytes (18). Given the lack of Ucp1 expression in hepatocytes, the increased EE (or oxygen consumption) in response to GCGR activation, is likely driving ATP synthesis. This raises the interesting question of which specific metabolic processes are operating at such high energy costs in the liver under chronically active GCGR signaling. One viable option are futile cycles (FCs), which are opposing chemical reactions that run simultaneously without generating a net product but waste ATP. Considering our data, two possible types of FCs could be contributing to the hypermetabolic state. First, futile lipid cycles constitute the continuous hydrolysis of triglycerides into glycerol and acyl-CoA units followed by the re-esterification into triglycerides. It is conceivable that glycerol and free fatty acids (FFA) derived from white adipose tissue (WAT) lipolysis are re-esterified in the liver and form a systemic futile lipid cycle, while the necessary ATP is delivered from the oxidation of FFA from the circulation and intrahepatic stores (19). The consequent efficient removal of FFA and glycerol from the periphery would further explain why potentially higher lipolytic rates in WAT of BI 456908-treated mice are not mirrored in elevated levels of these metabolites.

One of the major concerns for the clinical use of dual agonists with a GCGR-component is potential hyperglycemia. Hepatic GCGR stimulation promotes glucose output from the liver by increasing endogenous glucose synthesis and glycogen breakdown. Thus, liver specific ablation of GCGR leads to reduced fasting blood glucose levels compared to WT mice (20). Interestingly, chronic activation of GCGR by BI 456908 did not elevate fasting blood glucose levels. These observations introduce the second feasible futile cycle, which leverages the gluconeogenic capacity of the liver (21). The reciprocal pathways of gluconeogenesis and glycolysis harbor two reaction steps that can build futile cycles including the bi-directional conversions of glucose-6-phosphate to glucose and of fructose-1, 6-bisphosphate to fructose-6-phosphate. The resultant cycles might add to the preserved hypoglycaemia, suggesting that newly synthesized gluconeogenic intermediates are entrapped in the last steps of the pathway, entering futile cycles that could simultaneously modify EE. The concomitant activation of GCGR/GLP1R led to comparable improvements in glucose tolerance as single GLP1R agonism, independent of the genotype. These results favour a dominant glucose-regulatory function of the GLPR1R component as main effector. Indeed, survodutide is ineffective in ameliorating glucose tolerance in GLP1R knockout mice (2) and GLP1R agonism increases glucose-stimulated insulin secretion after two weeks of treatment, which enhances peripheral glucose disposal (22).

In conclusion, we demonstrate that hepatic GCGR activation is responsible and necessary for the substantial additive weight-loss effect of dual GCGR/GLP1R agonists, emphasizing the clinical relevance of GCGR agonism in the liver. The available literature and our data unambiguously converge on an overall mechanistic concept, suggesting that GCGR signaling in the liver drives mitochondrial fatty acid oxidation for ATP production, which might be fuelled by adipose tissue lipolysis and the utilisation of liver triglycerides, thereby increasing systemic energy expenditure.

## Materials and Methods

### Animal experiments

All animal procedures in this study were approved by the Cantonal Ethics Committee of the Veterinary Office of the Canton of Zurich. Male mice with a C57BL/6N background were used for the experiments. They are housed 2-5 littermates per cage in ventilated cages at standard housing conditions (22 °C, 12 h reversed light/dark cycle, dark phase starting at 7am, 40% humidity), with ad libitum access to high fat diet (HFD) (60% (kal%) fat, diet no. 3436, Provimi Kliba SA), and ad libitum access to water. At the end of the study, animals were euthanized individually by a CO2 overdose.

### Peptides Administration

The compounds were dissolved in a phosphate buffer (50 mM, pH 7.0) with 5% mannitol, which was also used as the vehicle control. All peptides were administered at a volume of 5 mL/kg. The first dose was given in the morning (dark cycle for animals) of Day 0.

### Hepatocyte-specific GCGR KO mouse model

GCGR floxed mice were generated by the Center for Transgenic Models, University of Basel, Switzerland. Hepatic-specific GCGR knockout (KO) mice were generated by crossing GCGR floxed mice with Albumin-Cre mice (#035593, JAX), resulting in Alb-GCGR fl/fl mice with liver-specific depletion of GCGR. These hepatic-specific GCGR KO mice, aged 6-8 weeks, were challenged with HFD for 16 weeks prior to compound treatment.

### Body composition measurement

Body composition measurements were performed on live mice using quantitative nuclear magnetic resonance imaging (EchoMRI). Fat and lean mass was analysed using the Echo MRI1 14 software.

### Intraperitoneal glucose tolerance tests

Intraperitoneal glucose tolerance test (ipGTT, 2g/kg BW) was performed after 21 days of treatment with mono or dual agonists after a four hours fasting period. Tail vein blood glucose concentrations were measured at 0, 15, 30, 60, and 120 minutes using a commercially available glucometer (ACCU-CHEK Aviva, Roche) with test strips. The area under the curve was calculated from glucose concentrations measured between 0 and 120 minutes.

### Plasma parameters

Blood samples were collected by cardiac puncture into EDTA-coated tubes from mice that were treated with compound and fasted for 4 hrs. Plasma was obtained by centrifugation at 8,000g/20 min/4°C. Plasma Glycerol and NEFA levels were determined using the Free Glycerol Reagent (Sigma-Aldrich) and NEFA assay kit (Wako NEFA kit), respectively. Plasma TAG levels were determined using the TAG assay kit (Roche Trig/GB reagent), and total cholesterol levels were measured with LabAssat™ Cholesterol kit (FUJIFILM). Plasma ALT activity was measured using the ALT activity assay (Sigma-Aldrich). All assays were performed according to the manufacturers’ instructions.

### Hepatic TAG level determination

To measure hepatic TAG levels, liver samples (50-100 mg) were weighed and homogenized in 1 ml of isopropanol per 50 mg tissue. The homogenates were incubated with rotation before centrifugation at 2,000g/10 min/4°C to collect the supernatant. TAG levels were determined using Roche Trig/GB reagent and normalized to the weight of the liver samples.

### Quantification and statistical analysis

Littermates were randomly assigned to treatment groups for all the experiments. Sample size was determined based on previous experiments. Animal numbers for each treatment are indicated in the corresponding figure legends. Data are represented as mean±SEM, unless it is indicated specifically. Data were statistically analysed by ANOVA using GraphPad Prism 10 software. The significance of treatment difference was analyzed using two-way ANOVA with Tukey’s post hoc multiple comparison tests with a *p<0.05 considered as significant.

## Acknowledgements

The authors meet criteria for authorship as recommended by the ICMJE. The authors did not receive payment related to the development of this publication. Boehringer Ingelheim was given the opportunity to review the manuscript for medical and scientific accuracy as well as intellectual property considerations. The study was supported and funded by Boehringer Ingelheim. BI 456908 is a back-up compound to survodutide and is licensed to Boehringer Ingelheim from Zealand Pharma.

